# Hsf1 is SUMOylated in the activated trimeric state

**DOI:** 10.1101/2020.10.05.327064

**Authors:** Szymon W. Kmiecik, Katarzyna Drzewicka, Frauke Melchior, Matthias P. Mayer

**Affiliations:** Center for Molecular Biology of Heidelberg University (ZMBH), DKFZ-ZMBH-Alliance, Im Neuenheimer Feld 282, D-69120 Heidelberg, Germany; Oncoarendi Therapeutics SA, 101 Zwirki i Wigury street, PL02089 Warsaw

**Keywords:** Protein misfolding, stress response, transcription regulation, Heat shock factor protein 1 (HSF1), small ubiquitin-like modifier (SUMO), post-translational modification (PTM)

## Abstract

The heat shock response (HSR) is a transcriptional program of organisms to counteract an imbalance in protein homeostasis. It is orchestrated in all eukaryotic cells by heat shock factor 1 (Hsf1). Despite very intensive research, the intricacies of the Hsf1 activation-attenuation cycle remain elusive at a molecular level. Posttranslational modifications belong to one of the key mechanisms proposed to adapt the Hsf1 activity to the needs of individual cells and phosphorylation of Hsf1 at multiple sites has attracted much attention. According to cell biological and proteomics data, Hsf1 is also modified by SUMO (small ubiquitin-like modifier) at several sites. How SUMOylation affects Hsf1 activity at a molecular level is still unclear. Here, we analyzed Hsf1 SUMOylation *in vitro* with purified components to address questions that could not be answered in cell culture models. *In vitro* Hsf1 is primarily conjugated at lysine 298 with a single SUMO, though we did detect low level SUMOylation at other sites. None of the tested E3 SUMO ligases increased SUMOylation efficacy as compared to the level in the presence of high concentrations of the E2 Ubc9. We provide evidence that Hsf1 trimerization and phosphorylation at serines 303 and 307 increases SUMOylation efficiency, suggesting that Hsf1 is SUMOylated in its activated state. Hsf1 can be SUMOylated when DNA-bound, and SUMOylation of Hsf1 does neither alter DNA binding affinity nor does it affect Hsc70 and DnaJB1-mediated monomerization of Hsf1 trimers and concomitant dislocation from DNA. We propose that SUMOylation acts at the transcription level of the HSR.

## Introduction

Protein misfolding is detrimental for cells due to loss of function but also due to gain of toxicity of some misfolded and aggregated protein species. To cope with proteotoxic stress a highly conserved homeostatic transcriptional program, the so-called heat shock response (HSR), emerged early in cellular evolution. The main regulator of the HSR in eukaryotic cells is the heat shock transcription factor Hsf1. Like with other transcription factors, the activity of Hsf1 is controlled on many levels (1, 2). In mammals, Hsf1 is in monomer-dimer equilibrium in unstressed cells (3). Upon proteotoxic stress Hsf1 trimers accumulate in the nucleus and bind to heat shock elements, three to four inverted NGAAN repeats, in promoters and enhancers to drive transcription of heat shock genes (4). One of the key mechanisms of Hsf1 regulation that allows the protein to respond to changing environmental or physiological conditions are posttranslational modifications (PTMs) (5).

Several studies focused on phosphorylation and acetylation of Hsf1 (4–10). Hsf1 phosphorylation at multiple sites is observed upon stress induction of the HSR and was considered a hallmark of Hsf1 activation (4, 11–14). Albeit, phosphorylation at most sites seem to downregulate Hsf1 transactivation, whereas only Hsf1 phosphorylation at S230 promoted Hsf1 activity (7). Similar to phosphorylation, Hsf1 acetylation can also up- and downregulate Hsf1 activity. Hsf1 acetylation at K80 was shown to reduce DNA binding of Hsf1 and was proposed to be important for attenuation of the HSR (9). Likewise, acetylation at K118 by E1A Binding Protein P300 (EP300) was proposed to impair functionality of Hsf1 (10). In contrast acetylation at K208 and K298 by EP300 reduced degradation of Hsf1 through the proteasome resulting in enhanced HSR (10).

Less studied is Hsf1’s post-translational modification with SUMO (small ubiquitin-like modifier), although SUMOylation per se has been intensively studied in the context of transcriptional regulation (15–17) and in heat shock response (see below). SUMOylation is a reversible and highly dynamic modification that regulates the function of more than thousand proteins, many of which are chromatin-associated (reviewed in (18–20)). SUMO specific conjugating enzymes (E1 activating, E2 conjugating and E3 ligating enzymes) form an isopeptide bond between SUMO’s Carboxy-terminus and the ε-amino group of lysines in an ATP-dependent reaction cascade, whereas SUMO isopeptidases revert the modification by hydrolysis. Many proteins that are SUMOylated carry a short SUMOylation consensus motif, ΨKxE (where **Ψ** is a large hydrophobic residue and **x** is any residue), that is recognized by the SUMO E2 conjugating enzyme Ubc9. Vertebrates express at least three SUMO proteins: SUMO2 and SUMO3 are virtually identical, frequently form chains via a SUMOylation consensus motif in their flexible N-termini, and their modification is strongly stimulated upon stress including heat shock. SUMO1, which shares only 50% sequence identity with SUMO2/3, exists largely in conjugated form under normal growth conditions and does usually not form chains as it lacks the N-terminal SUMOylation site (20–22).

The SUMO stress response (SSR) is activated upon heat stress and results in rapid conjugation of SUMO2/3 to protein substrates (16, 23, 24). This general protein SUMOylation is proposed to be an early reaction to protein misfolding, protecting partially misfolded proteins by increasing their solubility through addition of SUMO chains (25). Consistently, cells depleted for SUMO2/3 are more sensitive to heat stress (26).

The SSR also leads to massive changes in the proteome, suggesting an influence on gene expression. SUMOylated proteins are found at sites of actively transcribed inducible genes (like heat shock genes) (26-Lyst:2007ck 28). On one side, SUMO2/3 modification upon heat stress upregulates genes connected with survival, growth but also cell death (27). On the other side, SUMO2/3 modification upon heat stress represses mostly genes associated with transcription, reducing the overall load on the protein quality surveillance machinery (pro survival function). SUMOylation was proposed to inhibit transcription re-initiation, thereby sustaining Polymerase II (Pol II) pausing (24, 26, 28, 29). In acute stress, increase in SUMO modification is correlated with the occupancy of heat shock gene promoters by PIAS1 SUMO E3 ligase and RNA Pol II. Thereby, SUMOylation has been proposed as a mechanism to tightly regulate heat shock genes by preventing transcriptional hyperactivation (28).

How SUMO exerts its function in transcription regulation is largely unknown. On one side, stress-induced SUMO modification on active chromatin was proposed to act indirectly by stabilizing protein complexes on DNA rather than to act on transcription directly (27). On the other side, transcription factors are frequent targets for SUMO modification (15, 16).

Interestingly, Hsf1 is very rapidly and transiently SUMOylated upon heat stress (26, 28, 30). This transient modification is linked to phosphorylation in proximity to a conventional SUMOylation consensus motif surrounding K298, a finding which led to the first description of a phosphorylation-dependent SUMOylation motif (PDSM) (30, 31). The described PDSM consists of 8 residues: **ΨKxExxSP**, where **Ψ** is a large hydrophobic residue and **x** is any residue. Ser 303, which is part of the PDSM, and Ser 307 (Fig 1A) can both be phosphorylated in response to heat shock. S307 phosphorylation by MAPK may be required for modification of S303 by GS3K (13), but this is a matter of debate (30). Increased transcriptional activity has been observed for the non-SUMOylatable Hsf1-K298R and Hsf1-S303A variants in a cell culture model, strongly implying that SUMOylation downregulates the transactivation activity of Hsf1 (31). It has been proposed that this phosphorylation-dependent SUMOylation regulates Hsf1 by reducing its transactivation capability (30).

**Figure 1:**
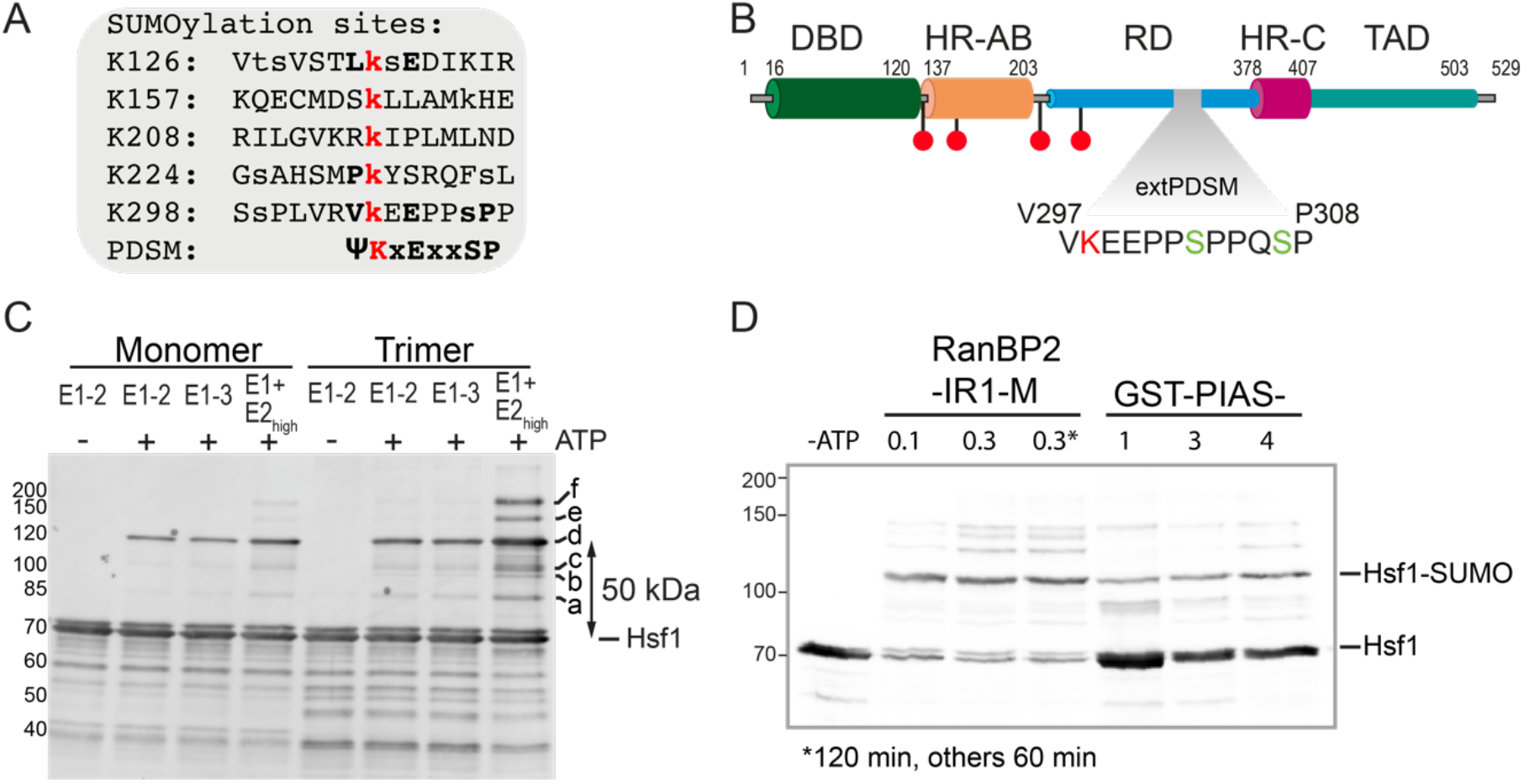
Hsf1 is efficiently SUMOylated *in vitro* by E1 and E2 enzymes. **A**, SUMOylation sites within Hsf1 identified in high through-put mass spectrometry studies. Modified lysins are indicated in red. Other potentially modified residues are indicated with lower cases. Residues consistent with the phosphorylation dependent SUMOylation motif (PDSM) are shown in bold. **B**, Hsf1 domain organization. The extended phosphorylation dependent SUMOylation motif (extPDSM) is indicated. Phosphorylated serines (S303 and S307) are indicated in green, SUMOylated lysine residue (K298) is indicated in red. The position of other lysine residues reported to be SUMOylated are indicated (red). **C**, Scanning for the best SUMOylation conditions for Hsf1 trimer and monomer. Monomeric and trimeric Hsf1-S303E,S307E were incubated with N-His-SUMO1 (10 μM), Aos1/Uba2 E1 enzyme (0.1 μM), Ubc9 E2 (low: 0.25 μM or high: 1.1 μM) in the presence and absence of the E3 SUMO ligase domain of RanBP2 (0.05 μM) and ATP (5 mM) as indicated, incubated for 2 h at 25°C and subsequently separated by SDS-PAGE, blotted onto a PVDF membrane and detected with an Hsf1-specific antiserum. Molecular weights in kDa are indicated on the left. **D**, SUMOylation efficiency screening in the presence of different E3 ligases (RanBP2-IR1-M, PIAS1, PIAS3, PIAS4). Trimeric Hsf1 wild type was SUMOylated in the presence of Aos1/Uba2 (0.2 μM), Ubc9 (0.5 μM), untagged SUMO2 (8 μM) and ATP (5 mM). RanBP2 fragment was tested at 0.1 and 0.3 μM concentrations, samples were analysed after 60 min of incubation or after 120 min of incubation (0.3*).

Whether K298 is the only relevant SUMOylation site in Hsf1, however, is not clear. Proteomics identified four additional SUMOylation sites in Hsf1 (K126, K157, K208, K224) (Fig 1B) (32, 33). One of these sites (K208) is within the sequence essential for monomerization and dislocation of Hsf1 from DNA by the Hsp70 machinery (34) and may affect this process. Consistent with the possibility that Hsf1 may carry more than one SUMO is an unusual low electrophoretic mobility of SUMOylated Hsf1 in SDS-PAGE (Hietakangas 2003).

Moreover, how SUMOylation regulates Hsf1 at a molecular level remained unclear in the published cell culture-based studies. Whether Hsf1 can be SUMOylated before activation in the monomeric state or subsequent to activation, in the trimeric state, and how exactly this modification influences Hsf1-DNA binding also remained unknown. The role of phosphorylation in regulating Hsf1 SUMOylation was also not clear. To address these questions, we reconstituted Hsf1 SUMOylation *in vitro* with purified components. We demonstrate that Hsf1 can be SUMOylated *in vitro* with high efficiency. Consistent with cell culture studies our data indicate that lysine 298 is the preferred SUMOylation site within Hsf1 *in vitro*. Furthermore, we show that most Hsf1 molecules are conjugated with a single SUMO, and we only find traces of double SUMOylated species, suggesting a low degree of SUMO chain formation or additional SUMOylation sites *in vitro*. Mass spectrometry and biochemical data strongly imply that Hsf1 mono SUMOylation results in a 50-kDa-upshift observed on SDS-PAGE, most likely due to branched conjugate formation. In our *in vitro* system trimeric Hsf1 is more efficiently SUMOylated than monomeric Hsf1. The phosphomimetic Hsf1-S303E,S307E variant is slightly better SUMOylated *in vitro* than wild type Hsf1, indicating that Hsf1 phosphorylation at S303/307 is not strictly required for SUMO transfer onto Hsf1. SUMOylation did not interfere with Hsf1-DNA binding or Hsc70-mediated dissociation of Hsf1 from DNA. Thus, we propose that Hsf1 SUMOylation attenuates Hsf1 action by interfering with the process of transcription itself, for example, by recruiting corepressors to heat shock gene promoters or by impairing the interaction of Hsf1 transactivation domain with the transcription machinery.

## Results

### In vitro Hsf1 is predominantly mono-SUMOylated at a single site

To clarify the nature of Hsf1 SUMOylation, we turned to *in vitro* SUMOylation experiments with full length Hsf1. In light of the observation that heat stress induced Hsf1 SUMOylation requires phosphorylation, we decided to test both wild type Hsf1 and phosphomimetic variant. As serines 303 and 307 are both phosphorylated under stress conditions, we considered the double phosphomimetic variant Hsf1 S303E, S307E to best imitate the *in-vivo*-situation.

In a first step, the best conditions for Hsf1 SUMOylation *in vitro* had to be established. The SUMOylation reaction is catalyzed by a system of three enzymes: the E1 SUMO activating enzyme (heterodimer Aos1/Uba2), the E2 conjugating enzyme (Ubc9) and an E3 ligase (like PIAS or RanBP2). *In vitro* a combination of E1 and E2 seems to be sufficient to transfer SUMO to the consensus site containing substrate and E3 ligases are proposed to accelerate the reaction or drive substrate specificity (18).

Preliminary screening for Hsf1 SUMOylation conditions demonstrated no increase of efficiency in the presence of E3 ligases (RanBP2−IR+M or PIAS, Fig1C and D), instead combination of E1 (0.1 μM) enzyme with the E2 enzyme at high concentration (1.1 μM) was sufficient to boost Hsf1 SUMOylation to high extent *in vitro* (Fig 1C).

One might wonder whether the obtained SUMOylation yield *in vitro* resembles the extent to which Hsf1 is SUMOylated under physiological conditions. According to published cell culture studies only minor fractions of Hsf1 seem to be SUMOylated (30, 31), as also observed for other transcription factors (22). Hsf1 *in vitro* SUMOylation yields in our study may be even higher than what is observed in cell culture lysates with overexpressed Hsf1 as estimated from the published immunoblots most likely due to the absence of SUMO-isopeptidases in our experiments.

Under the optimal conditions, we observed six bands above unmodified Hsf1 that are only visible in the presence of the SUMOylation machinery and ATP but not in the absence of ATP, with the dominant band (d) approximately 50 kDa above unmodified Hsf1. The N-His-SUMO1 that was used in these initial experiments has a theoretical mass of 13164.66 Da (measured 13164.33 Da) but according to previous reports mono-SUMOylation causes an increase in apparent molecular weight on SDS-PAGE of 15–17 kDa (35). Therefore, we were intrigued by the fact that the molecular weight shift for the dominant band (d) above unmodified Hsf1 was around 50 kDa according to SDS-PAGE, arguing against Hsf1 mono-SUMOylation (also observed in (30)). In addition, another study investigating Hsf1 SUMOylation reported upon Hsf1 SUMOylation a shift smaller than 50 kDa arousing controversies (36). Three explanations appear possible for the observed 50 kDa increase in molecular weight of modified Hsf1. (i) The branched Hsf1-SUMO conjugate has a strongly reduced electrophoretic mobility in SDS-PAGE; (ii) Hsf1 is simultaneously SUMOylated at more than one site; (iii) SUMO-chains are formed on Hsf1 that increase the molecular weight of the conjugate. To address this question, we analyzed Hsf1 SUMOylated in the monomeric and trimeric state by mass spectrometry (MS) after enrichment by size exclusion chromatography (SEC). To prevent spontaneous Hsf1 trimerization of monomerirc Hsf1 *in vitro* (3) in all experiments that contained monomeric Hsf1 SUMOylation was performed at 25°C; experiments that contained only trimeric Hsf1, SUMOylation was performed at 30°C to increase the yield. SEC showed SUMOylated Hsf1 in the trimeric as well as in the monomeric fractions (Fig 2A and B). The mass spectrum revealed that 52% of Hsf1 were unmodified, 45% were modified by a single SUMO and less than 3% by two SUMO molecules (Fig 2C). Comparing these values with a quantification of the bands of the immunoblot indicates that the modification with a single SUMO can cause a shift of up to 50 kDa in SDS-PAGE (Fig 1C and Fig 2C). The fainter bands (a-c) between the band of unmodified Hsf1 and the major SUMOylation band (d), are likely to also represent mono-SUMOylated Hsf1, albeit at different sites that cause a smaller electrophoretic mobility shift, indicating that the observed electrophoretic mobility shift is very sequence context dependent. According to the phosphor-site database (www.phosphosite.org October 2020) Hsf1 can be SUMOylated at 5 different lysines: K126, K157, K208, K224, and K298 (Fig 1A and B). The bands above the major SUMOylation band (bands e and f) then most likely represent modifications with two SUMO moieties (at different sites or formation of SUMO chains). In these preliminary experiments (Fig 1 and 2) N-terminally histidine-tagged SUMO1 was used in the SUMOylation reactions, however, as additional amino acids introduced to a protein may change its properties, untagged versions of SUMO1 and SUMO2 were used in all further experiments.

**Figure 2:**
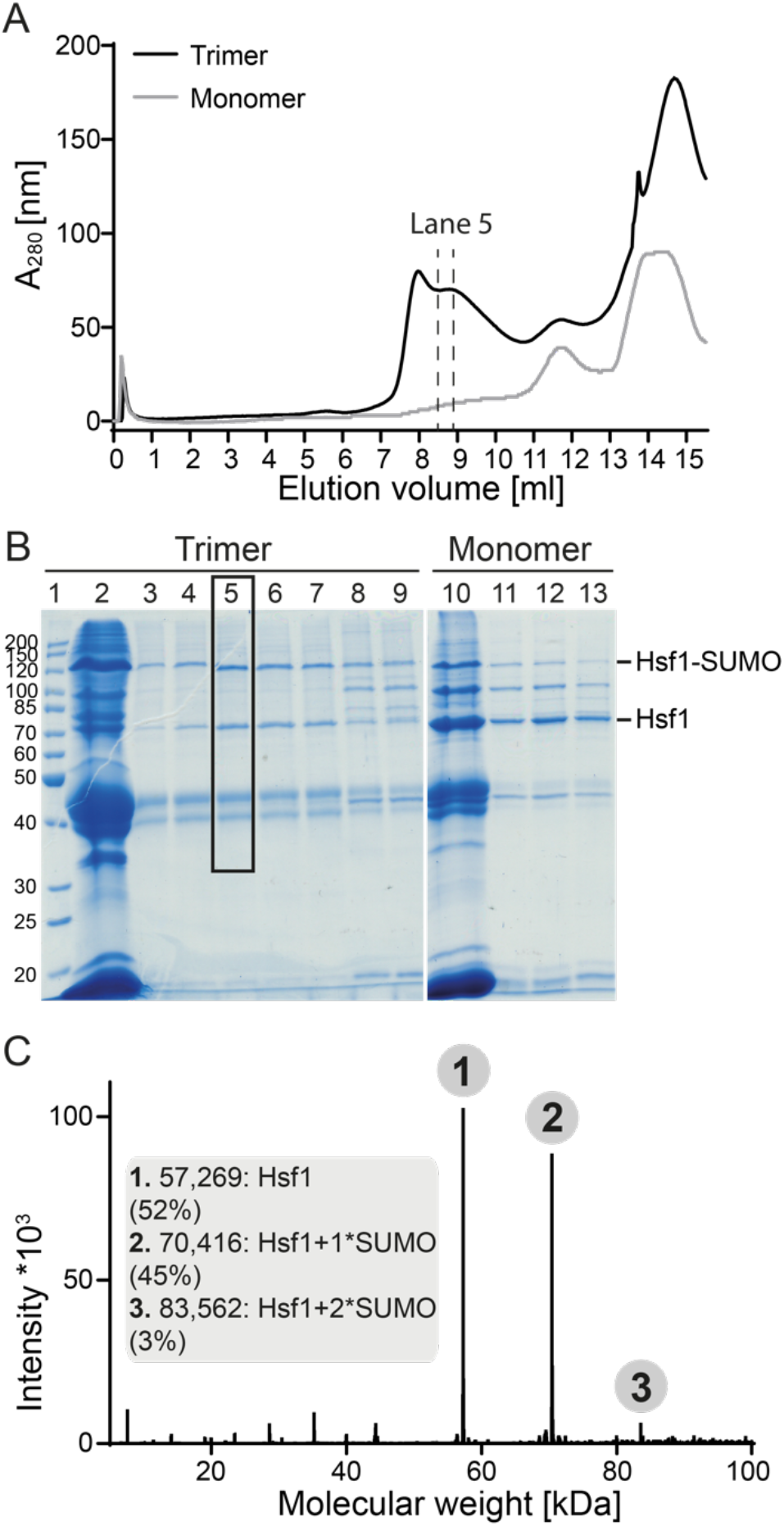
Hsf1 is mostly mono-SUMOylated *in vitro*. **A**, Size exclusion chromatography elution profile of the SUMOylation reaction for trimeric and monomeric Hsf1-S303E,S307E. Elution only to 15.5 ml is shown for clarity. Fraction analyzed by mass spectrometry (trimeric Hsf1 preparation) is indicated by dashed lines. **B**, SDS-PAGE analysis of collected fractions; Trimer: Lane 1: protein ladder; Lane 2: Load; Lanes 3-7: 0.5 ml fractions 7.42-9.92 ml; Lanes 8, 9: 0.5 ml fractions 10.92-11,92 ml. Fraction analyzed by mass spectrometry is indicated with the frame. Monomer: Lane 10: Load, Lanes 11-13: 0.5 ml fractions 10.92-12.42 ml. Molecular weights in kDa are indicated on the left. **C**, Deconvoluted MS spectrum of the analyzed fraction (Lane 5) corresponding to the mixture of SUMOylated and non-SUMOylated Hsf1-S303E,S307E. Detected molecular weights together with abundance percentage are indicated in the bracket. N-His-SUMO1 was used in this experiment. Theoretical mass of Hsf1-S303E,S307E alone is 57270.43 Da (measured 57269.30 Da). Theoretical mass of N-His-SUMO1 alone is 13164.66 Da (measured 13164.33 Da). Theoretical mass of mono-SUMOylated Hsf1 equals 70417.08 Da and for double-SUMOylated 83563.73 Da

### K298 is the primary Hsf1 SUMOylation site in vitro and its SUMOylation does not require phosphorylation

Demonstrating that Hsf1 can be mainly mono-SUMOylated, we wondered whether K298 is the major SUMOylation site within Hsf1 in our *in vitro* system. To evaluate this hypothesis *in vitro* K298 was replaced by arginine in the double phosphomimetic Hsf1-S303E,S307E variant. Consistent with the published *in vivo* data, the major band (d) of SUMO-Hsf1 almost completely disappeared upon replacement of K298 with arginine, but the bands a and c were still visible, arguing that these bands represent Hsf1 species that are SUMOylated at any of the other sites found by proteomics (Fig 3C and E). Phosphorylation on S303 was proposed to be an essential prerequisite for Hsf1 SUMOylation in cells (30). This is not the case *in vitro* where also wild type Hsf1 can be SUMOylated with SUMO2 albeit with lower yields than the phosphomimetic protein (Fig 3F). Phosphorylation within the SUMOylation motif was proposed to stabilize the interaction with the E2 modifying enzyme Ubc9 by providing additional electrostatic contacts (37).

**Figure 3:**
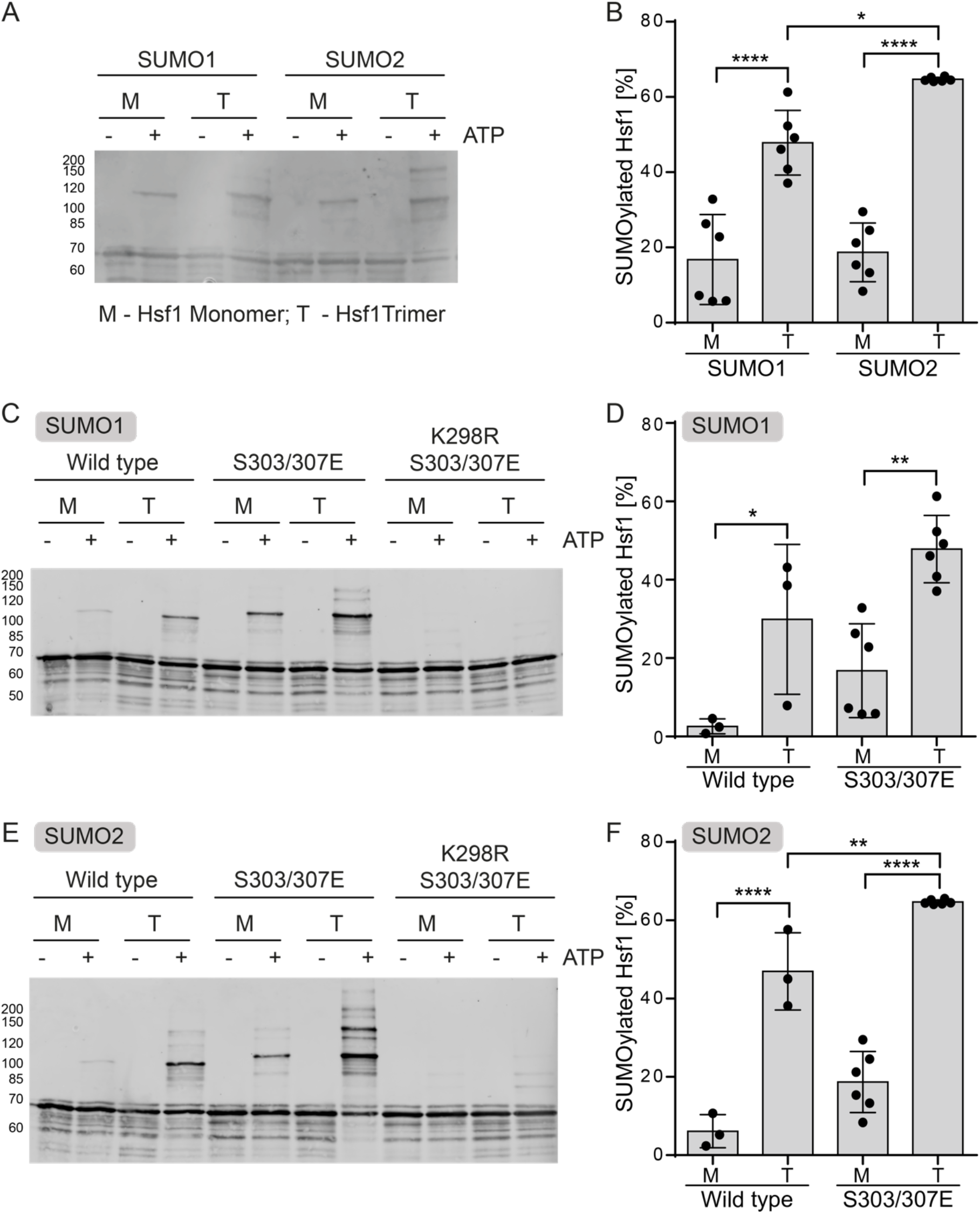
Lysine K298 is the primary SUMOylation site within Hsf1. **A**, Comparison of *in vitro* SUMOylation efficiency for SUMO1 (SMT3C) and SUMO2 (SMT3A). Hsf1-S303E,S307E was SUMOylated in the presence of Aos1/Uba2 (0.1 μM), Ubc9 (1.1 μM), untagged SUMO1 (10 μM) or SUMO2 (10 μM) and ATP (5 mM). Samples were prepared as indicated, separated by SDS-PAGE, blotted onto a PVDF membrane and detected with an Hsf1-specific antiserum. A representative Western blot is shown. Molecular weights in kDa are indicated on the left. **B**, Quantification of SUMOylation results from Figure 2A and similar immune blots. Percentage of mono-modified conjugate is calculated. Shown are mean ± SD, (n=6, ANOVA, *, p < 0.05; ****, p < 0.0001). **C**, Comparison of SUMOylation efficiency for Hsf1 wild type, Hsf1-S303E,S307E and Hsf1-K298R,S303E,S307E with SUMO1. Samples were prepared as indicated, separated by SDS-PAGE, blotted onto a PVDF membrane and detected with an Hsf1-specific antiserum. A representative Western blot is shown. Molecular weights in kDa are indicated on the left. **D**, Quantification of SUMOylation results from figure 2C. The percentage of mono-modified conjugate was calculated. Shown are mean ± SD, (wild type n=3, S303/307E n=6, ANOVA, *, p < 0.05; **, p < 0.01). **E**, Comparison of *in vitro* SUMOylation efficiency for wild type Hsf1, Hsf1-S303E,S307E and Hsf1-K298R,S303E,S307E with SUMO2. Samples were prepared as indicated, separated by SDS-PAGE, blotted onto a PVDF membrane and detected with an Hsf1-specific antiserum. A representative Western blot is shown. Molecular weights in kDa are indicated on the left. **F**, Quantification of SUMOylation results from figure 2E. The percentage of mono-modified conjugate was calculated. Shown are mean ± SD, (wild type n=3, S303/307E n=6, ANOVA, **, p < 0.01; ****, p < 0.0001).

### Trimeric Hsf1 is more efficiently SUMOylated than monomeric Hsf1

Upon heat shock, Hsf1 shifts from a monomer-dimer equilibrium to the trimeric state upon which it acquires DNA binding competency and released paused RNA-polymerase for transcription of heat shock genes. The heat shock response is attenuated by Hsp70-mediated monomerization and dissociation of Hsf1 from DNA. Evidence was provided that SUMOylation attenuates Hsf1 activity. There are several possible mechanisms how SUMOylation could influence Hsf1 driven heat shock gene transcription. (i) SUMOylation could prevent Hsf1 trimerization, (ii) SUMOylation could interfere with DNA binding or cause spontaneous dissociation from DNA (HSEs); (iii) Hsf1 SUMOylation could change the kinetics of Hsp70-mediated Hsf1 dissociation from DNA; (iv) the interaction with the transcription machinery could be modulated, transcription re-initiation may be inhibited after Hsf1 SUMOylation (as suggested in (28, 29)). The first scenario would imply that Hsf1 is SUMOylated in the monomeric state and that this modification slows down the transition to the trimeric DNA binding state. We therefore wondered whether Hsf1 is preferentially SUMOylated in the monomeric state or in monomeric and trimeric states to similar extent. To address this question, we generated Hsf1 trimers *in vitro* by heat shocking monomeric Hsf1 (10 min at 42°C). Our system allowed a direct comparison between SUMOylation yield for monomeric and trimeric Hsf1 *in vitro*. Hsf1 trimers were SUMOylated with higher (around 2 to 3-fold) efficiency than Hsf1 monomers for both SUMO1 and SUMO2 *in vitro* (Fig 3), suggesting that Hsf1 SUMOylation occurs subsequent to stress-induced trimerization and not as an initial response before Hsf1 trimerization and activation. Moreover, in our *in vitro* assays trimeric Hsf1 was more efficiently SUMOylated with SUMO2 (SMT3A) than with SUMO1 (SMT3C) (Fig 3A and B), consistent with the fact that upon proteotoxic stress increased SUMO2 but not SUMO1 modifications of Hsf1 were observed in cell culture studies (26, 31).

Since trimeric Hsf1 is more efficiently SUMOylated than monomeric Hsf1 (Fig 3) it is highly unlikely that Hsf1 SUMOylation interferes with trimer formation, excluding the first possibility.

### Hsf1 SUMOylation and Hsf1-DNA binding are independent and do not influence each other

To address the second possibility, we bound Hsf1 to fluorescently labeled DNA (HSEs) and subsequently added the SUMOylation machinery. Under these conditions about 25% of Hsf1 was SUMOylated (Fig 4B and C). However, we did not observe any significant decrease in fluorescence polarization during the SUMOylation reaction as compared to a reaction in the absence of ATP, which precludes SUMOylation, indicating that although Hsf1 is SUMOylated, it can still stay bound to DNA (HSEs) (Fig 4A). Moreover, an equilibrium titration to investigate the affinity of SUMOylated Hsf1 to DNA did not reveal any significant difference in K_D_ values (Fig 4D and E; Fig 4F indicates SUMOylation status of the analyzed samples). Consistent with Hietakangas and coworkers (30) our data demonstrate that binding of Hsf1 to DNA does not inhibit nor stimulate the SUMOylation process, implying that SUMOylation may occur after Hsf1 binding to DNA (Fig 4B and C). This observation, however, is not consistent with results of Hong et. al. showing that incubation of Hsf1, produced in reticulocyte lysate, with a semi-purified SUMO1 modification system increased binding of Hsf1wt but not Hsf1-K298R to DNA in an EMSA assay (36). Albeit, in this published experiment neither Hsf1 amounts nor SUMOylation of Hsf1wt was verified by immunoblotting.

**Figure 4:**
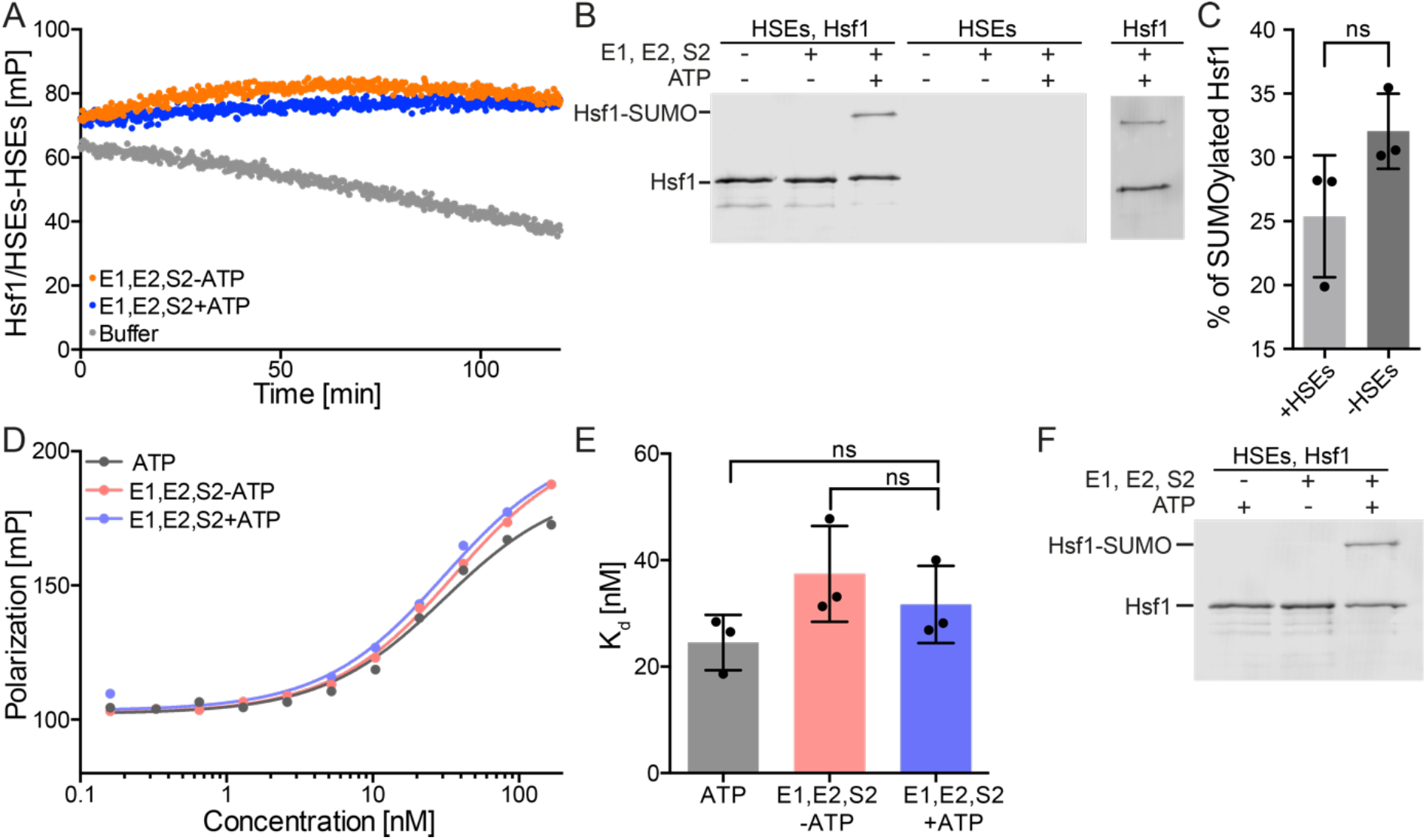
Hsf1 SUMOylation does not affect DNA binding and Hsp70-mediated Hsf1 dissociation from DNA. **A**, On DNA SUMOylation of Hsf1 monitored by fluorescence polarization. Hsf1-S303E,S307E was bound to fluorescent labeled HSE-DNA, then buffer or the SUMOylation machinery plus or minus ATP was added and fluorescence polarization monitored. Plotted are values corrected for control samples where Hsf1 was absent. **B**, Western blot of samples analyzed in Fig 4A. In addition to the right, a sample incubated in the absence of HSE-DNA with the SUMOylation machinery is shown. **C**, Quantification of SUMOylation efficiency in the presence and absence of DNA (HSEs). Shown are mean ± SD, (n=3, Student’s T-test, unpaired; ns, not significant). **D**, Equilibrium titration of Hsf1-S303E,S307E (SUMOylated trimeric Hsf1 versus non-SUMOylated samples) binding to Alexa 488-labeled HSE-DNA. Fluorescence polarization is plotted versus the Hsf1-S303E,S307E trimer concentration. **E**, DNA binding affinity for SUMOylated and non-SUMOylated Hsf1-S303E,S307E. 32% of Hsf1 was SUMOylated in this experiment (supplemental Fig. S3). Shown are mean ± SD, (n=3, ANOVA, Sidak’s multiple comparison; ns, not significant). **F**, A representative western blot of samples analyzed in panel **D**. 32% of Hsf1 was SUMOylated in this experiment.

To evaluate the influence of SUMOylation on Hsf1 dissociation from DNA (HSEs) SUMOylated Hsf1-S303E,S307E was subjected to Hsp70-mediated dissociation from DNA (HSEs). No significant differences in the kinetics of Hsp70-driven dissociation from DNA (HSEs) could be detected for the SUMOylated double phosphomimetic Hsf1 variant in comparison to the non-SUMOylatable Hsf1-S303E,S307E,K298R variant (Fig 5). The applied dissociation assay should be sensitive enough to detect differences in dissociation kinetics, if the SUMOylated Hsf1 fraction (28%, calculated in Fig 5C and D for time point 0) could not be dissociated from the DNA by the Hsp70 system (34). Moreover, Hsf1 SUMOylation did not impair the interaction with Hsp70 implying that *in vivo* Hsf1 de-SUMOylation by SUMO-isopeptidases could occur not only while Hsf1 is bound to DNA but also after Hsp70-mediated dissociation from DNA.

**Figure 5:**
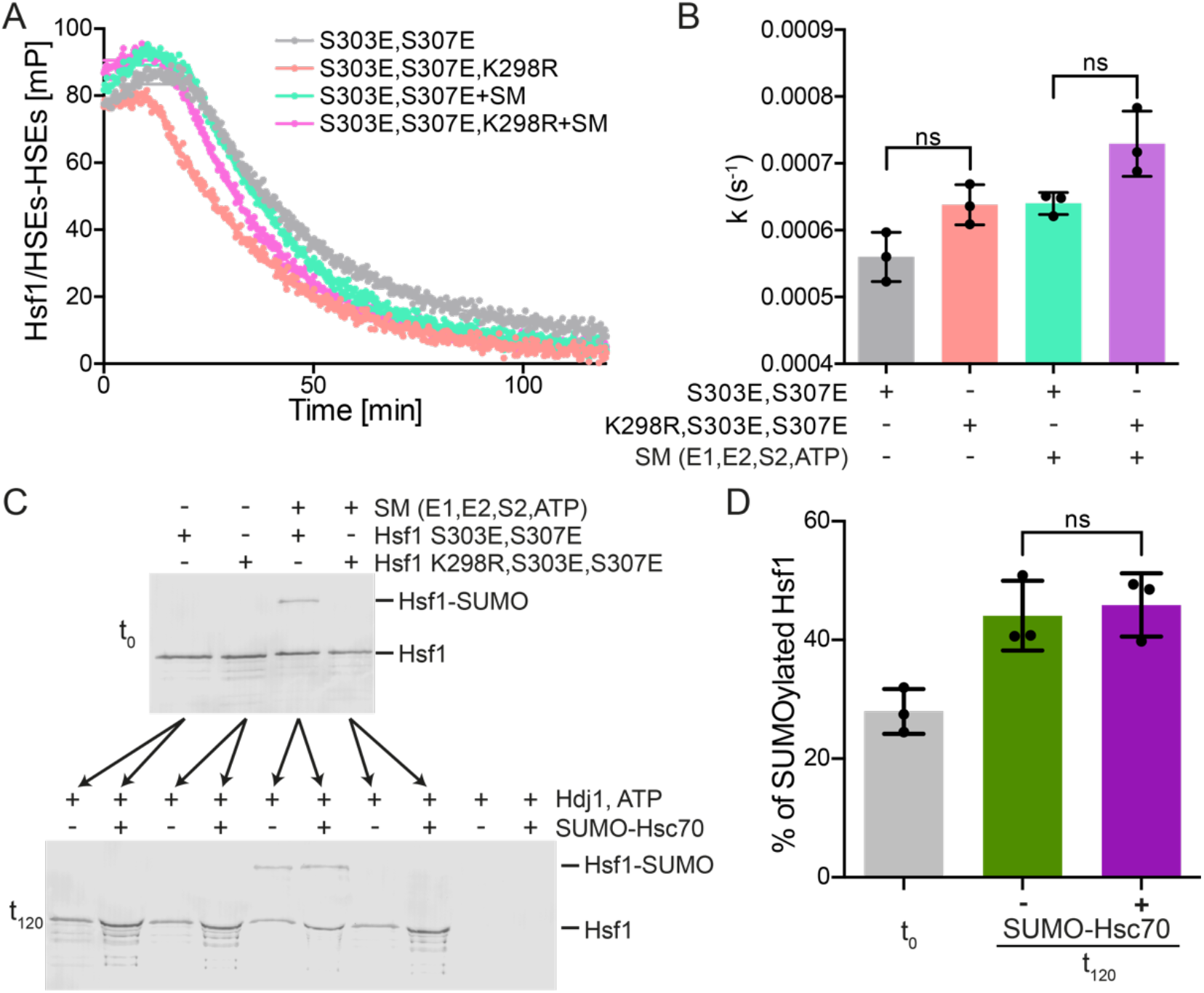
SUMOylation of Hsf1 does not influence Hsc70/DnaB1-mediated dissociation of Hsf1 from DNA. **A**, Dissociation of SUMOylated and non-SUMOylated Hsf1 from DNA (HSEs) by the Hsp70 system (SUMO-Hsc70, Hdj1/DNAJB1, ATP). Representative graph. **B**, Comparison of dissociation kinetics for different Hsf1 variants (SUMOylated S303E,307E and non-SUMOylable K298,S303E,S307E Hsf1 variants in the presence and absence of SUMOylation machinery). Shown are mean ± SD, (n=3, ANOVA, Sidak’s multiple comparison; ns, not significant). **C**, Representative western blot of samples before (t0) and after (t120) the dissociation kinetics measurement. **D**, Comparison of SUMOylation efficiency before dissociation reaction and after the reaction (sample in the presence and absence of SUMO-Hsc70). Shown are mean ± SD, (n=3,. ANOVA; ns, not significant).

## Discussion

In this study we answered several questions that could only be addressed in an *in vitro* reconstituted Hsf1 SUMOylation system with purified components. (1) We show that trimeric Hsf1 is SUMOylated more efficiently than monomeric Hsf1 suggesting that Hsf1 SUMOylation occurs after stress induced activation. (2) Hsf1 is primarily mono-SUMOylated and only a minor fraction carries two SUMOs. (3) Phosphomimetic Hsf1-S303E,S307E is SUMOylated only slightly better than Hsf1wt, demonstrating that SUMO conjugation does not strictly depend on prior phosphorylation. (4) SUMOylation of Hsf1 does not affect its DNA binding or its Hsc70/DnaJB1-mediated dissociation from DNA and monomerization.

In accordance with previous cell culture studies lysine 298 is the major SUMOylation site within Hsf1 *in vitro*, when using only the E1 SUMO activating enzyme Aos1/Uba2 and the E2 SUMO conjugating enzyme Ubc9, though we did observe some SUMOylation at other sites in Hsf1-S303E,S307E,K298R as well as in wild type Hsf1, consistent with proteomics studies (33). Therefore, our *in vitro* SUMOylation system mirrored the *in vivo* situation, though neither of the E3 ligases tested had a significant impact on the SUMOylation yield and the E1 and E2 enzyme sufficed *in vitro*. However, at low E2 concentrations and in the cellular context the situation might be different. Since Hsf1 is preferentially SUMOylated in the trimeric state and can be SUMOylated when bound to DNA, PIAS E3 ligases that are recruited to heat shock promoters may be able to enhance SUMOylation of Hsf1 due to high local concentrations or close proximity in the chromatin.

In our assays monomeric and trimeric Hsf1 can be distinguished, which is of advantage over previous cell culture studies where this was not feasible. We decisively demonstrate that trimerization and phosphorylation, two events accompanying Hsf1 activation, enhance Hsf1 SUMOylation *in vitro*. The crystal structure of Ubc9 in complex with RanGAP1 (37) reveals a positive patch at a position from the active center where phosphorylated S303 would be in Hsf1, suggesting that phosphorylation at S303 helps to position K298 in the catalytic center of Ubc9. Nevertheless, Hsf1 wild type protein can also be SUMOylated *in vitro*, indicating that S303 and S307 phosphorylation is not necessary for Hsf1 SUMOylation *in vitro*. This might also explain why the other potential SUMOylation sites are SUMOylated in our assay, albeit inefficiently, as they do not contain an acidic residue or a phosphorylatable serine or threonine five residues downstream of the SUMOylated lysine, a position corresponding to S303 for K298. Except for one (K126) they also do not contain a glutamate in position +2 or a large hydrophobic residue in position −1. The somewhat larger difference in SUMOylation efficiency between Hsf1wt and Hsf1-S303A *in vivo* as compared to the difference of unphosphorylated Hsf1wt and Hsf1-S303E,S307E *in vitro* may be due to several different effects. First, Hsf1-S303A may not be a good surrogate for unphosphorylated Hsf1, as the hydrophilic amino acid serine in position 303 may stabilize the Hsf1-Ubc9 complex more efficiently than the hydrophobic alanine. Second, Hsf1-S303E,S307E may not be a perfect surrogate for phosphorylated Hsf1, as the larger phosphate group might interact more favorably with Ubc9. Third, SUMOylated Hsf1wt might be more efficiently de-SUMOylated by SUMO-isopeptidases than phosphorylated Hsf1.

Hsf1 activity upon SUMOylation may be regulated by several different mechanisms. SUMO modification could compete with other modifications like acetylation or ubiquitination for target lysines (15, 22). Competition between SUMO and ubiquitin modification could lead to stabilization of the protein by preventing ubiquitin-mediated targeting to the proteasome (38–40). In such a case SUMOylation would be expected to increase Hsf1 concentrations and seems inconsistent with the inhibitory effect of SUMOylation on Hsf1 activity. Also, ubiquitination of K298 the major site for SUMOylation has not been reported so far, in contrast to ubiquitination of K208 that is likewise SUMOylated, albeit to much lower degree than K298. SUMOylation has been reported to affect the distribution of proteins between cytoplasm and nucleus (41). The subcellular localization of Hsf1 is currently still debated and some publications show Hsf1 mostly in the nucleus, whereas other publications show it in the cytoplasm in unstressed cells and in the nucleus after heat shock (42). Reduced import of Hsf1 into the nucleus could explain the SUMOylation-linked attenuation of the HSR. However, we consider such a mechanism for SUMO action on Hsf1 as less likely since our data show that Hsf1 is more efficiently SUMOylated in the trimeric state and Hsf1 is even SUMOylated when bound to DNA. SUMOylation has been described to inhibit DNA binding of the modified transcription factor as in the case of p53 (43), but also the contrary, to stimulate DNA binding as in the case of Hsf2 that is proposed to be converted into the DNA binding active state by modification with SUMO-1 at Lys82 in the DBD (44). Our study demonstrates that Hsf1 SUMOylation does not impair binding of this transcription factor to DNA (consistent with (31, 45)), excluding SUMOylation depended dissociation from DNA as mechanism for SUMOylation-induced attenuation of the HSR. In addition, binding of SUMO target proteins to DNA may regulate the modification efficiency *in vitro* and *in vivo* ((46) - PCNA, (47) - PARP-1, (48)). This is not the case for Hsf1 *in vitro*. DNA binding did not significantly increase Hsf1 SUMOylation yields under our conditions, indicating that SUMOylation and DNA binding are two independent events and this modification may occur before or after Hsf1 binding to DNA. Albeit, in the context of more extended DNA fragments or chromatin this might be different. Hsf1 SUMOylation also does not interfere with Hsp70-mediated dissociation of Hsf1 from DNA which has recently been proposed as a mechanism for Hsf1 activity attenuation (34). Our data suggest that Hsf1 SUMOylation very likely takes place after Hsf1 binding to DNA (in line with (24, 28)) and subsequent Hsf1 de-SUMOylation may occur after Hsp70-mediated Hsf1 dissociation from DNA. To regulate transcription Hsf1 needs to interact with a large number of factors including components of the transcriptional machinery and complexes remodeling the chromatin. Hsf1 SUMO modification can inhibit such interactions by masking an interaction interface or promote additional interactions for example with proteins containing a SUMO interacting motif (SIM), changing the interactome of Hsf1 (16, 17). SUMOylation has been described to promote or disrupt protein-protein interaction making the influence of this modification on gene expression context dependent (49). Thus, a possible explanation for SUMOylation driven Hsf1 activity attenuation could be that Hsf1 SUMOylation affects the interaction with the transcriptional machinery by modulating the interaction with positive transcription elongation factor (p-TEFb) (24) or by recruiting transcriptional co-repressors like histone deacetylases (HDACs) (17).

Taken together, our data supported by previous studies suggest sequential Hsf1 regulation model where stress induces Hsf1 trimerization, MAPK-mediated phosphorylation at serine 307 and GSK3-mediated phosphorylation of serine 303. Both events: trimerization and phosphorylation enhance Hsf1 SUMOylation efficiency. SUMO modification decreases Hsf1 activity by impairing interaction with activators of transcription or recruiting co-repressors of transcription. Both, modified and unmodified, forms of Hsf1 can be dissociated from DNA by the Hsp70 system to attenuate the HSR (Fig 6).

**Figure 6:**
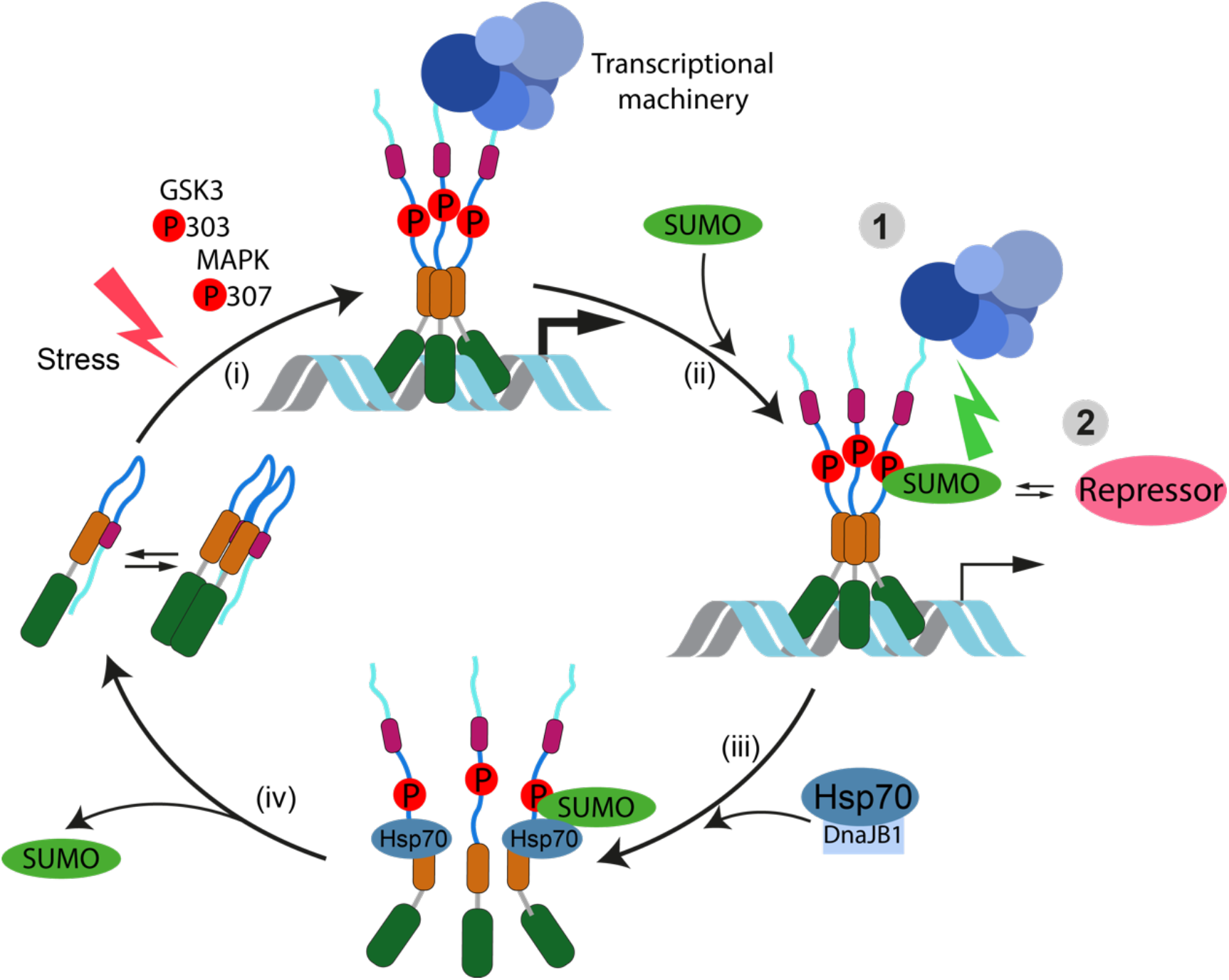
Model of Hsf1 activity regulation by SUMOylation. **(i)**, Hsf1 trimerizes upon stress. The activation is accompanied by Hsf1 phosphorylation. Among others serine 307 is phosphorylated by MAPK and serine 303 is phosphorylated by GSK3. **(ii)**, Upon transcriptional activation Hsf1 is SUMOylated by the SUMOylation machinery and its activity is repressed by 1: impairment of Hsf1 interaction with transcriptional machinery or/and 2: recruitment of co-repressors. **(iii)**, Hsf1-SUMOylation does not impair DNA binding or Hsp70-mediated dissociation from DNA. **(iv)**, Hsf1 SUMOylation is transient. Hsf1 is recovered to a resting state by the Hsp70 system.

## Experimental procedures

### Protein Expression and Purification

Human **Hsf1** was purified as a His6-SUMO fusion. *E. coli* BL21(DE3) Rosetta cells were freshly transformed with a plasmid encoding His6-SUMO-Hsf1. Overnight preculture was grown at 30°C then bacteria were grown at 37°C to an OD_600_ of 0.6. Temperature was shifted to 20°C and protein overproduction was induced by addition of 0.1 mM IPTG. The culture was grown for 2 h at 20°C and cells were subsequently harvested by centrifugation (5000 × g, 4°C for 10 min). Bacterial pellet was resuspended in 20 ml of Hsf1 lysis buffer (25 mM HEPES/KOH pH 7.5, 100 mM KCl, 5 mM MgCl_2_ and 10% glycerol). Bacterial suspension was drop-wise frozen in liquid nitrogen and stored at −80°C. All following steps need to be carried out at 4°C. Cells were disrupted using a Mixer Mill MM400 (Retsch, Haan, Germany) and resuspended in 200 ml Hsf1 lysis buffer supplemented with 3 mM β-mercaptoethanol, DNase and protease inhibitors (10 μg/ml aprotinin, 10 μg/ml leupeptin, 8 μg/ml pepstatin, 1 mM PMSF). The resulting lysate was centrifuged (33000 x g, 4°C for 45 min) to remove cell debris. The soluble fraction containing His_6_-SUMO tagged Hsf1 was incubated for 25 min at 4°C with 0.4 g Protino Ni-IDA resin (Macherey-Nagel, Düren, Germany) in a rotation shaker. The resin was transferred to a gravity-flow column and washed with 100 ml of Hsf1 lysis buffer. Bound protein was eluted with Hsf1 lysis buffer containing 250 mM imidazole, 3 mM β-mercaptoethanol and protease inhibitors. The His_6_-SUMO tag was cleaved off by incubation with TEV protease for 1 h at 4° C. The cleaved Hsf1 was further separated by size-exclusion on Superdex™200 HiLoad 16/60 column (GE Healthcare), equilibrated with Hsf1 SEC buffer (25 mM HEPES/NaOH pH 7.5, 150 mM NaCl, 10% glycerol, 2 mM DTT). The fractions containing either monomeric or trimeric Hsf1 were concentrated to 10 μM, flash-frozen in liquid nitrogen and stored at −80°C.

SUMOylated Hsf1 purification: After big scale SUMOylation the sample was subjected to size-exclusion on Superdex™200 10/300 GL column (GE Healthcare), equilibrated with Hsf1 SEC buffer (25 mM HEPES/NaOH pH 7.5, 150 mM NaCl, 10% glycerol, 2 mM DTT).

Human **Hsc70 (HSPA8)** was purified as a His_6_-SUMO fusion from overproducing *E. coli* BL21(DE3) Rosetta Cells were resuspended in Hsc70 lysis buffer (50 mM Tris pH 7.5, 300 mM NaCl, 5 mM MgCl_2_, Saccharose 10%) supplemented with 3 mM β-mercaptoethanol, DNase and protease inhibitors (10 μg/ml aprotinin, 10 μg/ml leupeptin, 8 μg/ml pepstatin, 1 mM PMSF). Bacteria were lysed using a chilled microfluidizer (MicroFluidizer EmulsiFelx-C5, Avestin, C505113) at a pressure of 1000 bar. The resulting lysate was centrifuged (33000 x g, 4°C for 45 min) and the supernatant was incubated for 25 min at 4°C with 2 g Protino Ni-IDA resin (Macherey-Nagel, Düren, Germany) in a rotation shaker. The resin was transferred to a gravity-flow column, washed with 200 ml of Hsc70 lysis buffer followed by high salt (Hsc70 lysis buffer but 1 M NaCl) and ATP (Hsc70 lysis buffer with 5 mM ATP) washes. Hsc70/Hsp70 were eluted with Hsc70 lysis buffer containing 300 mM imidazole, 3 mM β-mercaptoethanol and protease inhibitors. Fractions containing Hsc70 were desalted (HiPrep 26/10 Desalting column, GE Healthcare) to HKM150 buffer (25 mM HEPES/KOH pH 7.6, 150mM KCl, 5mM MgCl_2_) and digested with Ulp1 protease overnight. After proteolytic cleavage the protein was incubated with 1.5 g Protino Ni-IDA resin (Macherey-Nagel, Düren, Germany) for 25 min at 4°C to remove His_6_-SUMO (for purification of SUMO-Hsc70 overnight digestion with Ulp1 and second incubation with Protino Ni-IDA was omitted). Flow through was collected, desalted to HKM150 buffer, concentrated to 50 μM, aliquoted, flash-frozen in liquid nitrogen and stored at −80°C.

Human **DnaJB1/Hdj1** was purified without tag from *E. coli* BL21(DE3) Rosetta cells after overproduction for 4 h at 30°C. Cells were resuspended in DnaJB1 lysis buffer (50mM Tris/HCl, pH 8, 10mM DTT, 0.6% (w/v) Brij 58, 2 mM MgCl_2_) supplemented with DNase and 1 mM PMSF. Bacteria were lysed using chilled microfluidizer at a pressure of 1000 bar. The resulting lysate was centrifuged (33000 x g, 4°C for 45 min). One volume of buffer A (50mM sodium phosphate buffer pH 7, 5 mM DTT, 1 mM EDTA, 0.1% (w/v) Brij 58) was added to the clarified lysate. DnaJB1 was then precipitated with 65% ammonium sulfate. The obtained precipitate was diluted in buffer B (50 mM sodium phosphate buffer pH 7, 5 mM DTT, 1 mM EDTA, 0.1% (w/v) Brij 58, 2 M Urea) and dialyzed against buffer B. DnaJB1 was loaded onto a cation exchange resin (SP Sepharose) and eluted with a 0 to 666 mM KCl gradient in 15 CV. DnaJB1-containing fractions were combined, dialyzed against buffer C (50 mM Tris/HCl, pH 7.5, 2 M Urea, 0.1% (w/v) Brij 58, 5 mM DTT, 50 mM KCl) and subsequently loaded onto a hydroxyapatite resin. Bound protein was eluted with increasing concentration of buffer D (50 mM Tris/HCl, pH 7.5, 2 M Urea, 0.1% (w/v) Brij 58, 5 mM DTT, 50 mM KCl, 600 mM KH_2_PO_4_). DnaJB1-containing fractions were dialyzed against HKMG300 buffer (25 mM HEPES, pH 7.6, 5 mM MgCl_2_, 300 mM KCl, 10% glycerol), aliquoted, flash-frozen in liquid nitrogen and stored at −80°C.

### Purification of SUMOylation machinery components (see also (50))

Unless stated otherwise, protein purification protocols involved IPTG-induced expression in *E. coli* BL21 gold (Stratagene), bacterial lysis with lysozyme, and a 100,000xg spin for 1 h to collect soluble proteins. Each buffer contained 1 μg/ml each of leupeptin, pepstatin, and aprotinin, and 1 mM DTT (or β-mercaptoethanol); lysis buffers also contained 0.1 mM PMSF. After the specific purification steps described below, proteins were aliquoted, flash frozen, and stored at −80°C. The final buffer in each protocol was transport buffer (TB: 20 mM HEPES, 110 mM K-acetate, 2 mM Mg-acetate, 0.5 mM EGTA).

#### Human E1 enzyme

Purification involved co-expression of His-Aos1 and Uba2, bacterial lysis in 50 mM Na-phosphate (pH 8), 300 mM NaCl, 10 mM imidazol, purification on ProBond Resin (Invitrogen), size-exclusion on Superdex™200 HiLoad 16/60 column (GE Healthcare) and ion exchange chromatography (Mono Q, Pharmacia Biotech).

#### Human E2 enzyme (Ubc9)

Purification involved lysis in 50 mM Na-phosphate (pH 6.5), 50 mM NaCl, incubation of the 100,000xg supernatant with SP-sepharose beads (SIGMA), elution of Ubc9 from the beads with 50 mM Na-phosphate (pH 6.5), 300 mM NaCl, and size-exclusion on Superdex™200 HiLoad 16/60 column (GE Healthcare).

#### Human SUMO1/SUMO2/HisSUMO1

Purification involved bacterial lysis in 50 mM Tris/HCl (pH 8), 50 mM NaCl by sonification, preclearing of the 100,000xg supernatant with Q sepharose (SIGMA) in case of untagged SUMO (in case of HisSUMO1 nickel affinity chromatography was performed), concentration, and subsequent size-exclusion on Superdex™75 HiLoad 16/60 column (GE Healthcare).

### *In vitro* SUMOylation assays

#### Comparison of SUMOylation yield *in vitro*

Hsf1 *in vitro* SUMOylation reactions were set up to 20 μl in the assay buffer (20 mM HEPES/KOH pH 7.3, 110 mM KAcO, 2 mM Mg(AcO)_2_, 1 mM EGTA, 0.05% Tween, 0.2 mg/ml Ovalbumin, 1 mM DTT, 1 μg/ml of each aprotinin, leupeptin and pepstatin) and consisted of 0.1 μM SUMO E1 (His-Aos1/Uba2), 0.25 μM or 1.1 μM (high) SUMO E2 (untagged Ubc9), 50 nM GST-IR+M (fragment of RanBP2 E3 ligase, if applied), 10 μM His-SUMO1 or 9 μM SUMO1/SUMO2 (untagged), 0.5 μM of monomeric or trimeric (purified as a trimer or monomer heat shocked) Hsf1 (wt, S303E/S307E, K298R, S303E/S307E/K298R) and 5 mM ATP. After ATP addition, reactions were incubated for 3 hours at 25 °C. Samples were separated by SDS-PAGE, blotted onto a PVDF membrane and subsequently detected with an Hsf1-specific antiserum.

#### Using different E3 ligases

Hsf1 *in vitro* SUMOylation reactions were set up in the assay buffer (20 mM K-HEPES pH 7.3, 110 mM KAcO, 2 mM Mg(AcO)_2_, 1 mM EGTA, 0.05% Tween, 0.2 mg/ml Ovalbumin, 1 mM DTT, 1 μg/ml of each aprotinin, leupeptin and pepstatin) and consisted of 0.2 μM SUMO E1 (His-Aos1/Uba2), 0.5 μM SUMO E2 (untagged Ubc9), 0.1 or 0.3 μM GST-IR+M (fragment of RanBP2 E3 ligase) or 50 nM PIAS-1/-3/-4, 8 μM SUMO2 (untagged), 0.5 μM of trimeric Hsf1 wild type and 5 mM ATP. After ATP addition, reactions were incubated for h (2h for 0.3*) at 30 °C. Samples were separated by SDS-PAGE, blotted onto a PVDF membrane and subsequently detected with an Hsf1-specific antiserum.

### Fluorescence polarization assays

#### Preparation of fluorescently labeled DNA (ds-Alexa488-HSEs)

Fluorescently labelled ds-Alexa488-HSEs were prepared by annealing of fluorescently labelled Alexa488-3xHSE sense oligonucleotides (5’-[A488]-ccccTTCccGAAtaTTCcccc) with 3xHSE antisense nucleotides (5’-ggggGAAtaTTCggGAAgggg) (2 min at 70°C, 0.6°C/min stepwise decrease from 70°C to 30°C).

#### SUMOylation on DNA bound Hsf1

*In vitro* SUMOylation reactions were set up to 20 μl in the assay buffer (25 mM HEPES pH 7.6, 150 mM KCl, 5 mM MgCl_2_, 10% glycerol, 1 mM DTT) and consisted of 0.1 μM SUMO E1 (His-Aos1/Uba2), 1.1 μM SUMO E2 (untagged Ubc9), 10 μM SUMO2 (untagged), 5 mM ATP and 0.3 μM trimeric Hsf1 (S303/307E variant) bound to DNA (25 nM ds-Alexa488-HSEs). After samples preparation the plate was spun down at 1000 x g for 1 min at RT. Fluorescence anisotropy of the prepared samples was measured at 25°C for 2 h using a plate reader (CLARIOstar, BMG Labtech, Excitation, F:482-16, Emission, F:530-40). Samples were separated by SDS-PAGE, blotted onto a PVDF membrane and subsequently detected with an Hsf1-specific monoclonal antibody.

#### Binding of trimeric Hsf1 to Heat Shock Elements (HSEs)

5 nM ds-Alexa488-HSEs were titrated with trimeric Hsf1 at different concentrations ranging from 0.16 nM to 166.67 nM on low volume 384-well plate (CORNING, REF3820). SUMOylation was carried out at 30°C for 2 h (assay buffer: 25 mM HEPES pH 7.6, 150 mM KCl, 5 mM MgCl_2_, 10% glycerol, 1 mM DTT, E1/E2/ S2/Hsf1 equaled 0.08/0.92/8.33/1 μM and 4.17 mM ATP). The plate was subsequently spun down at 1000 x g for 1 min at RT. Fluorescence anisotropy of the prepared samples was measured after 15 min at 25°C using a plate reader (CLARIOstar, BMG Labtech, Excitation, F:482-16, Emission, F:530-40). The data points were fitted to the quadratic solution of the law of mass action using the GraphPad Prism software.

#### Hsf1 dissociation from Heat Shock Elements (HSEs) ((34) modified)

7.5 μM SUMO-Hsc70 and mM ATP in HKMG150 buffer (25 mM HEPES pH 7.6, 150 mM KCl, 5 mM MgCl_2_, 10% glycerol, 2 mM DTT) preincubated at 25°C for 30 min were mixed with 10 μM Hdj1/DNAJB1, 2 mM ATP, 100 nM trimeric Hsf1 and 25 nM HSEs preincubated at 25°C for 5 min on low volume 384-well plate (CORNING, REF3820) in a final 20 μl reaction volume. The plate was subsequently spun down at 1000 x g for 1 min at RT. Fluorescence anisotropy of the prepared samples was monitored in a plate reader (CLARIOstar, BMG Labtech, Excitation, F:482-16, Emission, F:530-40) at 25°C. A trimeric Hsf1 fraction was used for the experiments, if not stated otherwise in the figure legend. Data were normalized to samples where Hsf1 was absent. A single exponential equation with dissociation delay was fitted to the data points using the GraphPad Prism software:

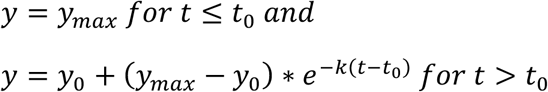

with *y*_*max*_ and *y*_0_ representing the fitted maximal and minimal fluorescence polarization values and *k* being the rate of the dissociation reaction.

##### SDS-PAGE

Protein samples pre-mixed with 5x SDS-PAGE sample buffer (250 mM Tris, pH 6.8, 50% glycerol, 10% SDS, 25% β-mercaptoethanol, 0.5% Bromophenol Blue) were separated on 12% SDS-PAGE gels in the Laemmli system (Running buffer: 25 mM Tris, 192 mM glycine, 0.1% SDS). Separation was carried out at constant current (50 mA per gel, 28 min). Gels were stained using a commercial staining solution (Quick Coomassie Stain, SERVA) or used further for Western blot analysis.

##### Western blot α-Hsf1

Proteins separated on PAGE gel were transferred to a PVDF membrane (Immobilon-P/Immobilon-FL, 0.45 μm, Merck Millipore) using Trans-Blot^®^ Turbo™ Transfer System (Bio-Rad). The membrane was subsequently blocked in PBS-T buffer (137 mM NaCl, 2.7 mM KCl, 10 mM Na_2_HPO_4_, 1.8 mM KH_2_PO_4_, 0.05% Tween 20) containing 1% milk for 45 min at RT. After blocking, the membrane was incubated with primary antibody (HSF1 (H-311) rabbit polyclonal IgG or HSF1 (E-4) mouse monoclonal IgG, Santa Cruz Biotechnology, 1:1000 dilution) overnight at 4°C on a roller. The next day the membrane was washed 3 times for 30 min with PBS-T buffer and incubated with the secondary fluorescently labelled antibody (Goat anti rabbit IgG IRDye 680RD or Goat anti-Mouse IRDye 800CW, Odyssay, 1:20000 dilution). After 1 h of incubation the membrane was washed 3 times for 30 min with PBS-T buffer. The protein of interest was further detected using fluorescence (LICOR Odyssey Infrared Imaging System, 700 nm or 800 nm channel). Western blots were quantified using Image Studio Lite Ver 5.2.

## Data availability

All data contained within the manuscript.

## Acknowledgements

We thank Dr. T. Ruppert and N. Lübbehusen for help in the core facility for mass spectrometry and proteomics, S. Hennes and A. Müller for excellent technical assistance. We kindly acknowledge Dr. Annette Flotho, Dr. Ramkumar Seenivasan and Dr. Roman Beloshistov for purified proteins and advise, and are particularly grateful to Dr. A. Werner for pilot experiments of Hsf1 *in vitro* SUMOylation. We also thank Dr. A. Wentink for suggestion to use SUMO-Hsc70 for dissociation experiments.

## Funding and additional information

This work was funded by the Deutsche Forschungsgemeinschaft (DFG, German Research Foundation) – Project-ID 201348542 - SFB 1036 TP9 to M.P.M. and SFB1036 TP15 to F.M.

## QUANTIFICATION AND STATISTICAL ANALYSIS

All biochemical assays were performed at least 3 times independently. Data were analyzed with GraphPad Prism 6.0 (GraphPad Software). Statistical significance was estimated by ANOVA or T-tests as indicated in figure legends. For Quantification ImageJ or Image Studio Lite Ver 5.2 were applied.

## COMMENTS

Please note, the nomenclature for mammalian SUMO2 and SUMO3 is used inconsistently. Like many colleagues in the SUMO field, we follow the nomenclature as introduced by Saitoh and Hinchey (23). Their assignment was consistent with the original description of mammalian SUMO genes (reviewed in (51)). According to this, mature SUMO2 (Smt3A) is 92 amino acids long, mature SUMO3 (Smt3B) consists of 93 amino acids.

